# Gene essentiality in cancer cell lines is modified by the sex chromosomes

**DOI:** 10.1101/2021.11.04.467330

**Authors:** Shahar Shohat, Ethel Vol, Sagiv Shifman

**Author notes:** Correspondence: Shahar Shohat, Department of Genetics, The Institute of Life Sciences, The Hebrew University of Jerusalem, Edmond J. Safra Campus, Jerusalem 91904, Israel., Sagiv Shifman, Department of Genetics, The Institute of Life Sciences, The Hebrew University of Jerusalem, Edmond J. Safra Campus, Jerusalem 91904, Israel.

## Abstract

Human sex differences are thought to arise from gonadal hormones and genes on the sex chromosomes. Here we studied how sex and the sex chromosomes can modulate the outcome of mutations across the genome. We used the results of genome-wide CRISPR-based screens on 306 female and 396 male cancer cell lines to detect differences in gene essentiality between the sexes. By exploiting the tendency of cancer cells to lose or gain sex chromosomes, we were able to dissect the contribution of the Y and X chromosomes to variable gene essentiality. Using this approach, we identified 178 differentially essential genes that depend on the biological sex or the sex chromosomes. Integration with sex bias in gene expression and the rate of somatic mutations in human tumors highlighted genes that escape from X-inactivation, cancer-testis antigens, and Y-linked paralogs as central to the functional genetic differences between males and females.

## Introduction

Males and females differ in many ways, among them the frequency of diseases (Cook et al., 2011; Cyranowski et al., 2000; Edgren et al., 2012; May et al., 2019; Ngo et al., 2014), the exhibition of symptoms for the same disease (Baba et al., 2005; Goldstein, 2006), and the response to different drugs (Wang et al., 2016). For example, in cancer, the frequency of most non-reproductive cancers is higher in males (Cook et al., 2011; Tevfik Dorak and Karpuzoglu, 2012). Some treatments have been shown to work differently in females and males for tumors with the same genetic characteristics (Pal and Hurria, 2010). The two main biological mechanisms that cause human sex differences are gonadal hormone secretions and genes located on the X and Y chromosomes, which we refer to as sex-specific genetic effects. There is considerable evidence for sex differences in diseases caused by gonadal hormone secretions (Baron-Cohen et al., 2005; Fuseini and Newcomb, 2017; Law et al., 2014). However, some sex differences are caused directly by sex-specific genetic effects (Baron-Cohen et al., 2005; Chen et al., 2012; Wang et al., 2016).

Sex-specific genetic effects stem from the genotypic difference between males and females: an XY genotype for males versus an XX for females. Substantial evidence exists for direct effects of the Y chromosome (Carruth et al., 2002; Charchar et al., 2012; Kido and Lau, 2015; Loke et al., 2015) and the X chromosome (Dunford et al., 2017; Kaneko and Li, 2018; Pal and Hurria, 2010; Snell and Turner, 2018) on human development and different diseases. The Y chromosome could affect sex differences through genes located at the male-specific region unique to the Y chromosome. Regarding the X chromosome, males have one X chromosome, and females have two. Still, most X chromosome genes do not show a significant difference in their expression level due to the X-inactivation process, which silences one of the X chromosomes. However, some genes escape the inactivation and are expressed from both copies. Those genes may have higher expression in females than males and contribute to the genetic sex differences (Disteche, 2012). Genes that escape X-inactivation are involved in cancer (Dunford et al., 2017; Kaneko and Li, 2018), developmental disorders (Adam and Hudgins, 2005; Peng et al., 2015; Snijders Blok et al., 2015), and autoimmune diseases (Syrett and Anguera, 2019; Youness et al., 2021).

Despite the evidence for sex differences in human diseases and the role of the sex-chromosomes in those diseases, the biological mechanisms that underlie sex differences are not fully understood. One approach for identifying sex-specific mechanisms is to explore genes with different functional importance in males and females, such as those essential more to one of the sexes. Recent advances in CRISPR technology have enabled the systematic identification of genes necessary for the normal functioning of cells across the genome (Shohat and Shifman, 2019; Tzelepis et al., 2016; Yilmaz et al., 2018). The Achilles project is the largest survey of human gene essentiality (Meyers et al., 2017). The project used CRISPR loss-of-function genome-wide screens to quantify the degree of gene essentiality (measured as gene dependency) in around 800 human cancer cell lines.

In this study, we use data from the Achilles project to identify genes more essential in one of the sexes. We harness the tendency of cancer cells to lose or gain sex chromosomes (Bianchi, 2009; Duijf et al., 2013; Kang et al., 2015a; Pageau et al., 2007; Richardson et al., 2006) to detect genes for which essentiality is dependent on the Y or X chromosomes. Our results reveal several mechanisms which underlie sex-dependent essentiality in cancer cells and the involvement of those processes in the sex-biased distribution of somatic mutation in cancer tumors.

## Results

### Identification of sex-dependent essential genes

To detect genes that show significant differences in their functional importance between the sexes, we compared the degree of gene essentiality between male and female cell lines. Essentiality measured as a dependency score (0 is equivalent to a gene under no selection and -1 is the median score of the common essential genes) was available for 18,119 genes in 306 female and 396 male cell lines. We identified 31 genes with significant (false discovery rate (*FDR*) < 0.1) sex-dependent essentiality (Table S1).

The cell lines originated from 29 different lineages, and among them are 111 cell lines from four tissues that only exist in females (female-only tissues: cervix, uterus, breast, and ovary). To ensure that the female-only tissues do not influence our analysis, we re-ran the analysis without the cell lines from the female-only tissues. Male-only tissues included only two cell lines, thus unlikely to affect the analysis. Two genes (CDK6 and PAX8) were no longer differentially essential (nominal *P > 0*.*05*) and therefore were excluded (Figure S1). Out of the remaining 29 genes, 24 were more essential to females (11 X-linked and 13 autosomal genes), and five were more essential to males (three X-linked and two autosomal genes) (Table S1, Figure 1A).

**Figure 1.**
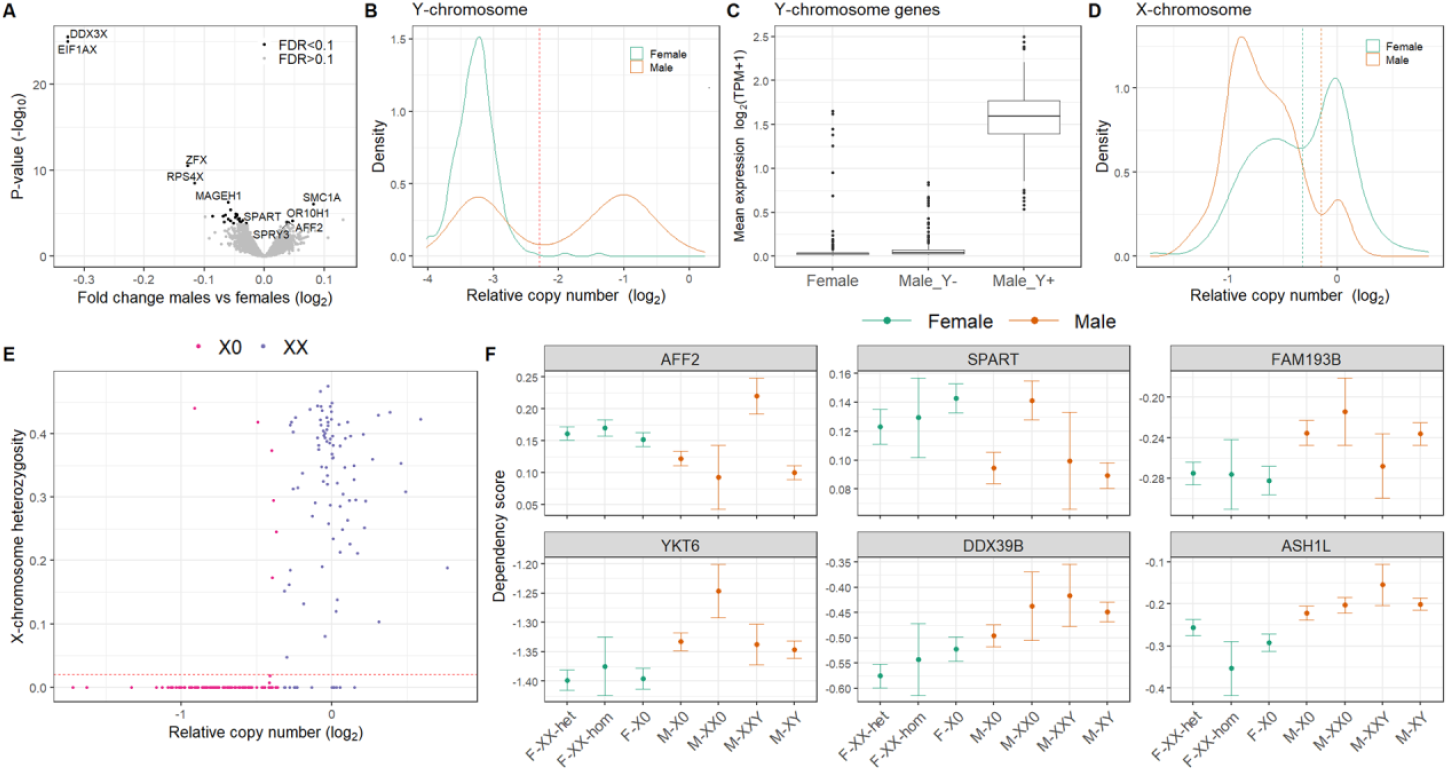
Identification of sex-dependent essential genes and sex chromosome ploidy. (A) Results of the differential essentiality analysis between male and female cell lines. Genes with a fold change < 0 are more essential to females, and genes with fold change > 0 are more essential to males. The labels are the names of the top five most significant genes that are more essential to each of the sexes. (B) Distribution of the mean relative copy number of Y-linked genes for male and female cell lines. The dashed line indicates the threshold for the definition as Y^+^ or Y^-^ cells. (C) The mean gene expression of Y chromosome genes for Y^+^ and Y^-^ cell lines, as inferred from the Y chromosome relative copy number. (D) Distribution of the mean relative copy number of X chromosome genes for male and female cells. The orange dashed line indicates the threshold used for definition as XX or X male cells, and the turquoise line shows the same for female cells. (E) The proportion of heterozygote SNPs as a function of the mean relative copy number (log2) of X chromosome genes in female cells. The dashed line indicates the threshold for classification as heterozygote or homozygote X chromosome. (F) Essentiality (dependency score) of the six most significant genes influenced only by biological sex. Values are the mean dependency score per genotype. Error bars are the standard error of the mean (SEM).

Human sex differences originate from a combination of genetic sex and gonadal hormone secretions. To separate the two causes, we leveraged the tendency of cancer cells to lose and gain sex chromosomes. To infer the number of Y chromosomes, we utilized the expression and copy number of Y chromosome genes (relative to all other genes) (Table S2). Classification based on the Y chromosome copy number (Figure 1B) was in high agreement with the classification based on the expression of Y chromosome genes (Figure 1C and S2A). Male cell lines with intermediate Y chromosome gene expression or females with high Y chromosome gene expression were excluded from further analysis (Figure S2A).

The number of X chromosomes was inferred from the relative copy number of X chromosome genes (Figure 1D). We also used the heterozygosity level of the X chromosome for female cells with available single nucleotide polymorphisms (SNPs) data (∼70%) (Figure 1E). Classification based on heterozygosity revealed that 94% of X0 female cells were homozygote for X chromosome SNPs, but this was also true for 15% of the XX female cells (Figure 1E). Heterozygote X0 female cells (n = 6) were removed from further analysis. Homozygote XX female cells (n = 15) were treated as a separate group in our study (XX_hom_). Loss of heterozygosity on the X chromosome is known to occur in cancer lines as a result of losing the inactive copy followed by a duplication of the active copy (Benoît et al., 2007; Kang et al., 2015a; Kawakami et al., 2004; Pageau et al., 2007; Richardson et al., 2006; Sirchia et al., 2005). We used the methylation level on the X chromosome as an indication for X-inactivation. As expected, X chromosome methylation levels were significantly higher in heterozygote XX females cell lines than other cell lines, including homozygote XX females (Figure S2B and Table S2).

After inferring the sex chromosome composition of the cells (Table 1), we estimated the separate effects of the sex chromosomes and the biological sex on the essentiality of the 29 sex-dependent genes (Table S3). Out of the 29 genes, seven were significantly (*P* < 0.05) associated with the Y chromosome, ten with the X chromosome, 20 genes were significantly associated with the biological sex, and six genes were significantly associated with more than one factor. The 15 genes influenced solely by the biological sex included 13 more essential to female cells and two more essential to male cells (Figure 1F and Figure S2C).

**Table 1.** Number of cell lines with different sets of sex chromosomes

| Genotype | Sample size |
| --- | --- |
| XX Heterozygote Female | 143 |
| XX Homozygote Female | 15 |
| XY Male | 181 |
| X0 Female | 134 |
| X0 Male | 142 |
| XX Male | 18 |
| XXY Male | 18 |

### The dosage of the sex chromosomes impacts gene essentiality

Our analysis shows that the presence or dosage of the sex chromosomes can explain a considerable proportion of the sex-dependent essentiality - a phenomenon we term sex-chromosome mediated essentiality (SME). We, therefore, decided to carry out a genome-wide screen for SME genes (SMEGs), studying the X and Y chromosomes separately.

To identify genes affected by the dosage of the X chromosome (X-SMEGs), we compared the essentiality levels of genes between XX heterozygote female (XX_het_) cells and X0 cells (both male and female). We identified 39 X-SMEGs (*FDR* < 0.1), of which 29 are X-linked genes (Figure 2A and Table S4). Only six of X-SMEGs were identified in the previous analysis of sex-dependent essential genes.

**Figure 2.**
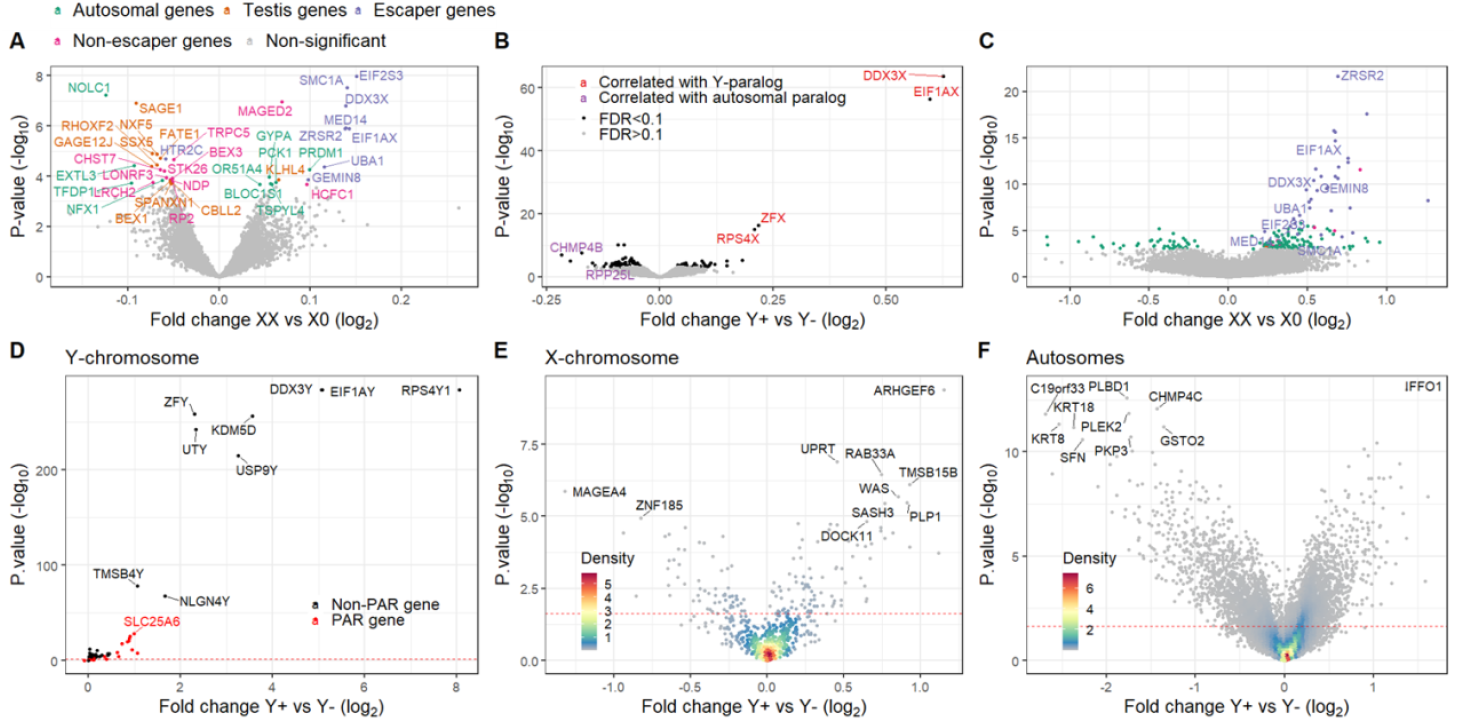
The effect of the sex chromosomes on gene essentiality and expression. (A) Differential essentiality analysis between XX_het_ and X0 cells. Genes with negative fold change (log_2_) are more essential to XX_het_ cells, and genes with positive value are more essential to X0 cells. The color of the gene names indicates different characteristics of the genes. (B) Differential essentiality analysis between Y^+^ and Y^-^ cells. Genes with a fold change < 0 are more essential for Y^+^ cells, and genes with fold change > 0 are more essential for Y^-^ cells. Labels are the names of genes likely to be explained by the expression of their paralogs. (C) Differential expression analysis between XX_het_ and X0 cells. Labeled genes are both differentially expressed (*FDR* < 0.1) and differently essential (*FDR* < 0.1) between XX_het_ and X0 cells. (D-F) Differential expression analysis between Y^+^ and Y^-^ cells. Results are shown separately for (D) the Y chromosome, (E) X chromosome, and (F) autosomes. Labels are the gene names of the top ten most significant genes in each category

To identify genes affected by the presence of the Y chromosome (Y-SMEGs), we compared gene essentiality between cells carrying a Y chromosome (Y^+^ male cells) and cells that do not (all female and Y^-^ male cells). The analysis was controlled for the effect of the number of X chromosomes and sex. We found 122 Y-SMEGs (*FDR* < 0.1). Among them, 42 genes were more essential to Y^-^ cells, and 80 were more essential to Y^+^ cells (Figure 2B and Table S5).

### Widespread effect of sex chromosome dosage on gene expression

Our discovery of SMEGs suggested that sex chromosome dosage could have general genetic effects. To investigate this further, we determined the impact on gene expression of X chromosome dosage and the presence of the Y chromosome.

To explore the effect of X chromosomes dosage, we performed differential expression analysis between XX_het_ and X0 cells. We found 183 significant genes (*FDR* < 0.1) (Table S5). As expected, the most significant differentially expressed genes were X-linked genes that escape from X-inactivation and are more expressed in XX compared to X0 cells (Figure 2C and Table S6). In total, 44 X-linked genes and 139 autosomal genes were differentially expressed (Figure 2B and Table S5).

Next, we studied the effect of the Y chromosome on gene expression by performing differential gene expression analysis between Y^+^ and Y^-^ cells (Figure 2D-F, Table S7). To our surprise, we detected 4,425 differentially expressed genes associated with the Y chromosome (*FDR* < 0.1). A large majority of those genes remained significant (85% with *FDR* < 0.1; 97% *P* < 0.05) when the analysis was restricted to male cells (XY vs. X0), as well as when XY male cells were compared to XX female cells (40% with *FDR* < 0.1; 56% *P* < 0.05) (Table S7). As expected, the biggest difference was observed for Y-linked genes and genes in the pseudoautosomal regions (PAR), including 36 Y-linked genes (80% of Y chromosome genes) and 13 PAR genes (70% of the PAR genes). Additionally, 150 X chromosome genes (not in the PAR) and 4,226 autosomal genes were differentially expressed.

### X-inactivation and testis-specific genes are involved in X-SMEGs

We wanted to characterize the X-SMEGs and determine possible mechanisms for the effect of the X chromosome on essentiality. Among the X-SMEGs, many are X-linked genes, but it is unclear why having two X chromosomes should modify essentiality if one is inactivated. We, therefore, tested if the X-SMEGs are enriched with genes that escape X-inactivation (Figure 2A). We found that X-SMEGs that are more essential to X0 than XX_het_ cells are indeed enriched for known escape genes (8 out of 17 genes; *P* = 1.5×10^−5^; odds ratio (OR) = 17.4). The eight X-SMEGs that escape from X-inactivation are significantly more expressed in XX_het_ cells (*FDR* < 0.05) and are highly essential across all cells (mean dependency score < -0.5). In contrast, out of the 22 X-SMEGs more essential to XX_het_ cells, 18 are X-linked genes, but only two escape from X-inactivation (Figure 2A, Table S4), and none are differentially expressed between XX_het_ and X0 cells (Figure 2C).

To characterize the 18 X-linked genes more essential to XX_het_ cells, we tested if they are preferentially expressed in any particular tissue. Interestingly, we found that those 18 X-linked genes are significantly enriched with genes selectively expressed in the testis (*P* = 9.3×10^−4^, FDR = 0.023) (Figure 3A). Nine of the 18 genes are expressed preferentially in the testis (CBLL2, GAGE12J, SAGE1, RHOXF2, BEX1, FATE1, SSX5 NXF5, and SPANXN1). Most of the testis-specific genes belong to a group known as cancer-testis antigens (CTAs). They are lowly expressed in almost all cells types outside the testis but show sporadic expression in cancer cells (Jungbluth et al., 2000; Simpson et al., 2005).

**Figure 3.**
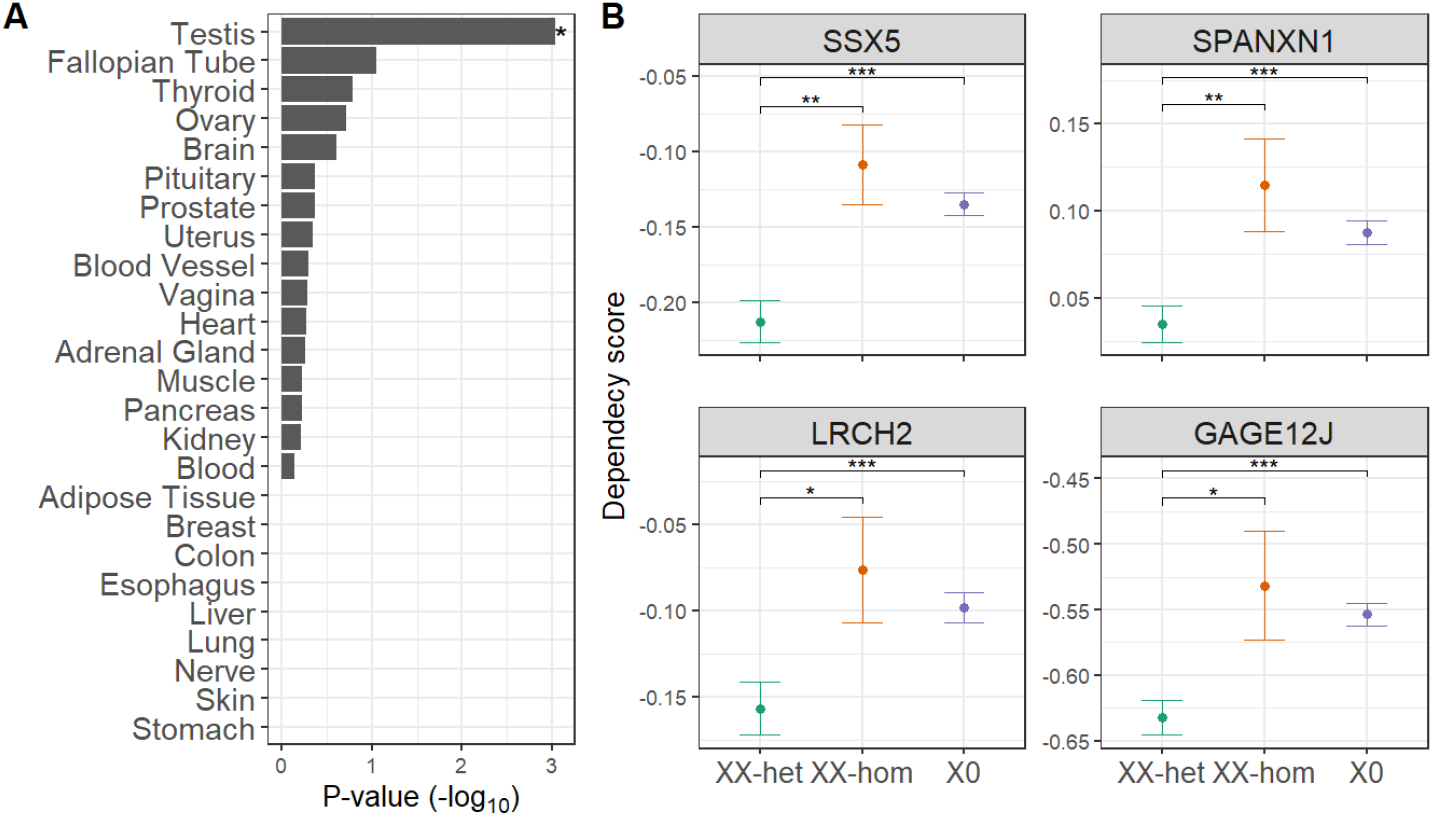
Testis-specific genes are more essential to XX_het_ cells. (A) X-linked genes that are more essential to XX_het_ cells are expressed preferentially in the testis but not in other tissues. ^*^ FDR corrected P < 0.05. (B) The top four most significant genes with higher essentiality to XX_het_ cells than X0 or XX_hom_ cells. ^***^ *P* < 0.001, ^**^ *P* < 0.01, ^*^ *P* < 0.05.

The reason why some genes are more essential to XX_het_ cells is not apparent. XX_het_ cells differ from X0 cells not only in X chromosome dosage but also in the inactivated X chromosome that exist only in XX_het_ cells. To determine if X-inactivation could be involved in the observed effects, we compared essentiality between XX_het_ cells to cells with two identical X chromosomes (XX_hom_ female and male cells). Based on the methylation levels, the XX_hom_ cells, despite having two X chromosomes, do not have an inactive copy (Figure S2B). We found that seven out of the 18 X-linked genes more essential to XX_het_ cells, including five CTA genes, are significantly more essential in XX_het_ than XX_hom_ cells (Figure 3B and Figure S3). It suggests that female cells with X-inactivation are more dependent on those testis-specific genes.

### Direct and indirect effects of the Y chromosome on gene essentiality

The top four significant Y-SMEGs are X-linked genes with a paralog on the Y chromosome. We assumed that the Y-linked paralogs might compensate for the loss-of-function mutations in the X-linked genes. We, therefore, tested the effect of paralog expression on Y-SMEGs. Out of the 122 Y-SMEGs, 29 genes had a paralog that was differently expressed between Y^+^ and Y^-^ cells (*FDR* < 0.1), and the expression of that paralog was significantly correlated with the levels of essentiality (*FDR* < 0.1) (Table S5). For six out of those genes, the correlation with the paralog was at the top two most significant correlations compared to all other genes in the genome (Figure 2B and Table S5). Four of the genes are X-linked genes (EIF1AX, DDX3X, RPS4X, and ZFX) with Y chromosome paralogs (EIF1AY, DDX3Y, RPS4Y1, and ZFY) (Figure 4A-C). The other two genes are autosomal (RPP25L and CHMP4B) with autosomal paralogs (RPP25 and CHMP4C) (Figure 4D-F). The two autosomal paralogs are significantly differentially expressed between Y^+^ and Y^-^ cells. These suggest that the presence of the Y chromosome can influence the essentiality of genes in two ways. The first is by a direct effect of Y-linked paralogs that can compensate for the loss of X-linked genes. Second, by indirectly influencing the expression of paralogs on the autosomes that can compensate for the loss of Y-SMEGs.

**Figure 4.**
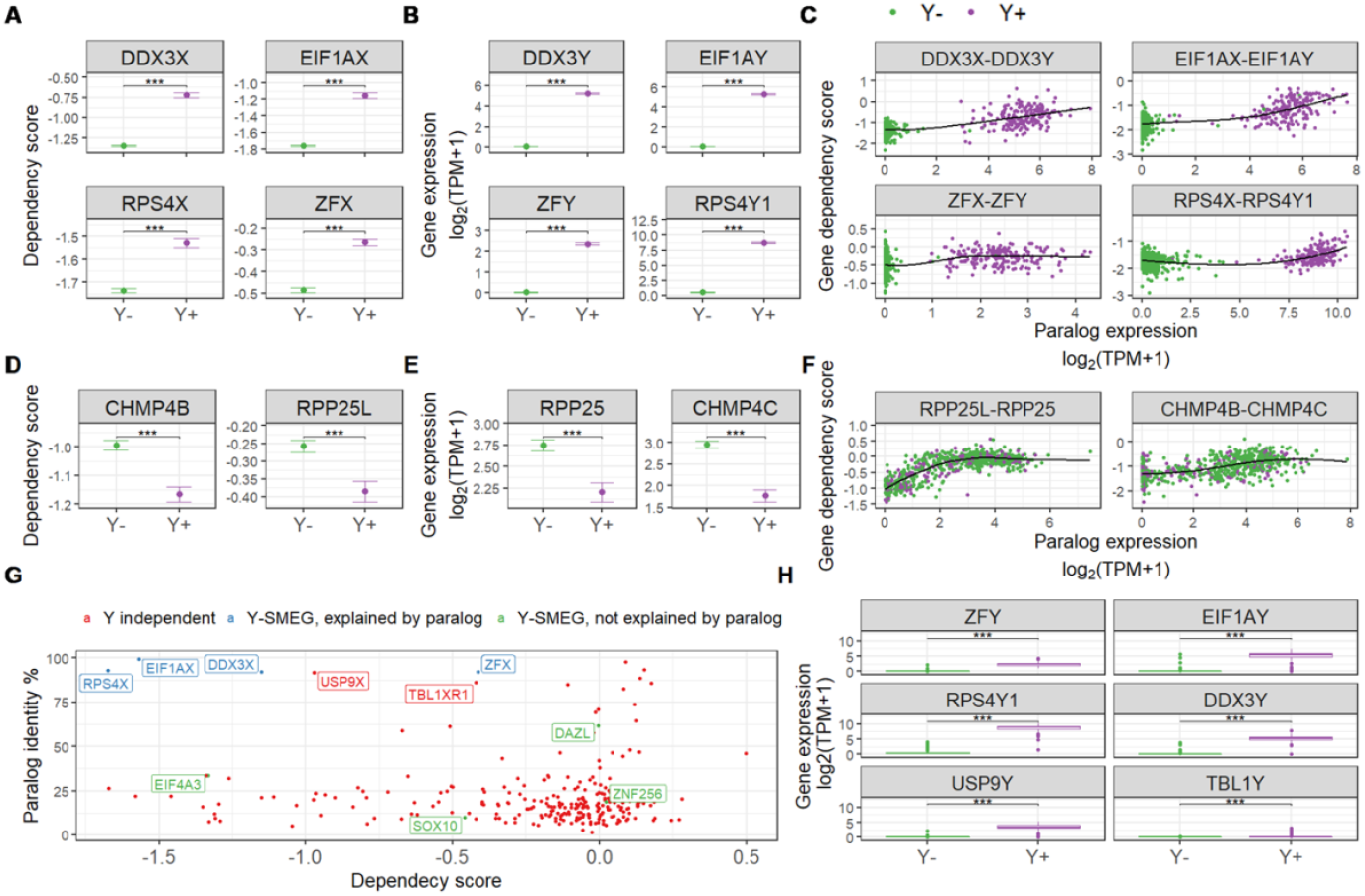
Paralog genes on both the Y chromosome and the autosomes can compensate for the loss of Y-SMEGs. (A-F) Evidence for six Y-SMEGs that can be compensated for their loss by a Y-dependent paralog. (A-C) For four X-linked genes, the paralog resides on the Y chromosome. (D-F) For two autosomal genes, the Y chromosome influences the expression of an autosomal paralog. (A, D) A significant difference in essentiality (dependency score) between Y^+^ and Y^-^ cells. (B, E) The expression of the paralogs in Y^+^ and Y^-^ cells parallel the differences in essentially. (C, F) Essentiality (dependency score) is significantly correlated with the gene expression of the paralog. The name of the Y-SMEG is written on the left and the Y-dependent paralog on the right. (G) The relationship between the sequence identity of the paralogs (percentage) and essentiality (mean gene dependency score), shown for all genes with a Y-linked paralog. The color of the gene names indicates Y-SMEGs explained (blue) or not explained (green) by a Y-linked paralog. In red label are two genes (USP9X and TBL1XR1) expected to be influenced by the Y chromosome because they have relatively high sequence identity with a Y-linked paralog and low dependency score. (H) The expression in Y^+^ and Y^-^ cells of six Y-linked genes that are paralogs of X-linked essential genes with high sequence identity between the paralogs. ^***^, *P* < 0.001.

The observation that four genes on the Y chromosome can compensate for the loss of their X-linked paralog raises the question of why other genes with paralogs on the Y chromosome are not Y-SMEGs. The four X-Y paralogs pairs we identified are characterized by a high similarity between the paralogs (> 80% sequence identity) and by being highly essential (mean dependency score across all cells = -1.2) (Figure 4G). In contrast, the 274 genes we identified with Y-linked paralogs tend to be either not very essential (mean dependency score = -0.19) or with low similarity to their Y-linked paralog (Figure 4G). We detected only two genes (USP9X and TBL1XR1) with high similarity to their Y chromosome paralogs (>85% sequence identity) that are relatively essential (mean dependency score = -0.98 and - 0.42, respectively) (Figure 4G). Regarding TBL1XR1, the reason for the lack of compensation may be the extremely low expression of the paralog gene (TBL1Y) in the majority of Y^+^ cells (Figure 4H). In contrast, the paralog of USP9X (USP9Y) is expressed in most Y^+^ cells (Figure 4H); thus, the lack of compensation ability may stem from functional divergence.

### Fine mapping of Y chromosome regions responsible for the differential essentiality

We found several mechanisms that may explain the Y-SMEGs; however, most genes remained unexplained. To map regions on chromosome Y associated with Y-SMEGs, we analyzed 14 cell lines with partial deletions of the Y chromosome (Figure 5A and Table S8). We used this approach to analyze all the Y-SMEGs and show the results for 31 genes with significant effects (Figure 5B-C and Figure S4-S5). Our previous analysis pointed to a highly likely causal gene in four out of the 31 genes (DDX3X, EIF1AX, ZFX, and RPS4X) (Figure 4A-C). The predicted section in the mapping analysis contained the causal gene (Figure 5C-C) for all four genes, supporting our approach.

**Figure 5.**
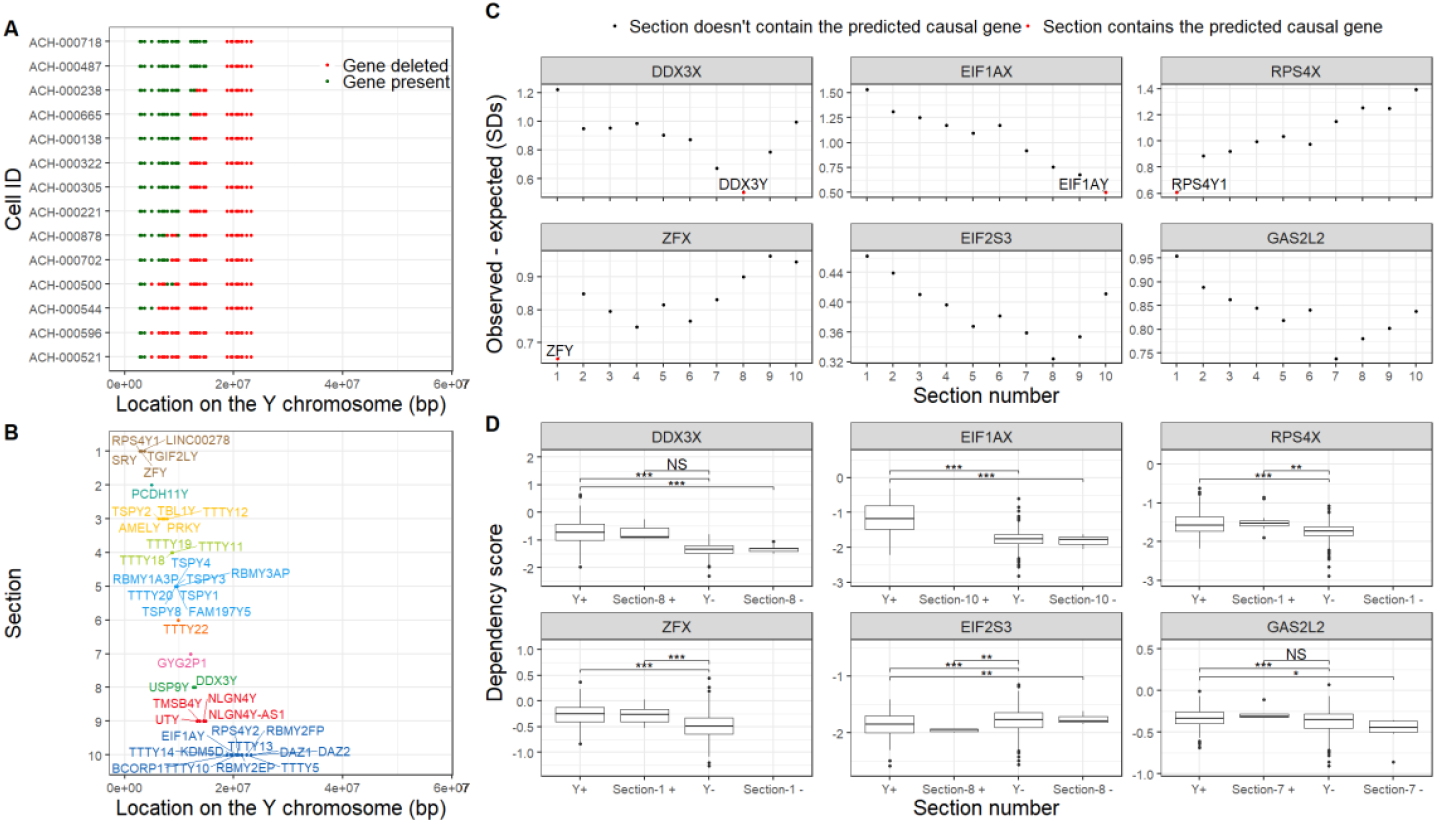
Partial deletions in the Y chromosome allow for fine mapping of regions responsible for Y-SMEGs. (A) A map of partial Y chromosome deletions in 14 cell lines. Each point is a gene with available copy number data, where green means that the gene exists and red means that it is deleted. (B) The names and location of Y-linked genes (with available copy number data), divided into ten sections. (C) The difference between the expected and observed dependency scores assuming the causal gene is in the indicated area. Values are the number of standard deviations. The results are shown for four genes with a suggested causal gene (DDX3X, EIF1AX, ZFX, and RPS4X) and two other genes with unknown causal genes. The labels are the names of the proposed causal genes. (D) Dependency score differences between Y^+^ and Y^-^ cells and between cells with and without the region most likely to include the causal gene. NS, *P* > 0.05; ^*^, *P*<0.05; ^**^, *P*<0.01; ^***^, *P*<0.001.

### Analysis of somatic mutations in cancer identifies genes with sex-biased mutation rates

We identified the effect of the sex chromosomes on gene essentiality in cancer cell lines. To test how these results relate to sex-dependent selection *in vivo*, we analyzed the sex-specific distribution of somatic mutations in cancer tumors. We used the Catalogue of Somatic Mutations in Cancer (COSMIC) (Tate et al., 2019), including somatic mutations in tumor samples from 4,755 females and 7,489 males collected from 12 different tissues. A previous analysis of a smaller, partially overlapping dataset (Dunford et al., 2017) identified six genes with sex-biased mutation rates. It is important to note that while the CRISPR screen in the cancer cell lines often results in a complete knockout of genes, most somatic mutations are in a heterozygote state with unknown functional consequences.

We compared the rate of nonsynonymous mutations between male and female tumors, and we identified 30 genes with significant differences (*FDR* < 0.1) (Figure 5A). All the identified genes are located on the X chromosome and none on the autosomes or the PAR regions. Three out of six genes previously identified (Dunford et al., 2017) were among the four most significant genes (KDM6A, DDX3X, and KDM5C). For 17 of the genes, the mutation rate was higher in females (female-biased genes), and for 13 genes, it was higher in males (male-biased genes) (Figure 6A, Table S9). None of the identified genes showed a significant difference in the mutation rate of synonymous mutations (Figure 6B, Table S11). Out of the 30 genes, four also showed differential essentiality depending on sex or sex chromosomes in the cancer cell lines (DDX3X, HTR2C, KLHL4, and THOC2). Two of those genes are highly essential (DDX3X and THOC2, average dependency score < -1.0), and the results agree with an increased mutation rate in the sex where the gene is less essential. In the case of DDX3X, which is more essential to females and Y^-^ cells, there are significantly more nonsynonymous mutations in male tumors (male: female (M: F) ratio = 2.3; *FDR* = 0.00061), while THOC2 that is more essential to Y^+^ cells shows significantly more nonsynonymous mutations in female tumors (M: F ratio = 0.61; *FDR* = 0.018).

**Figure 6.**
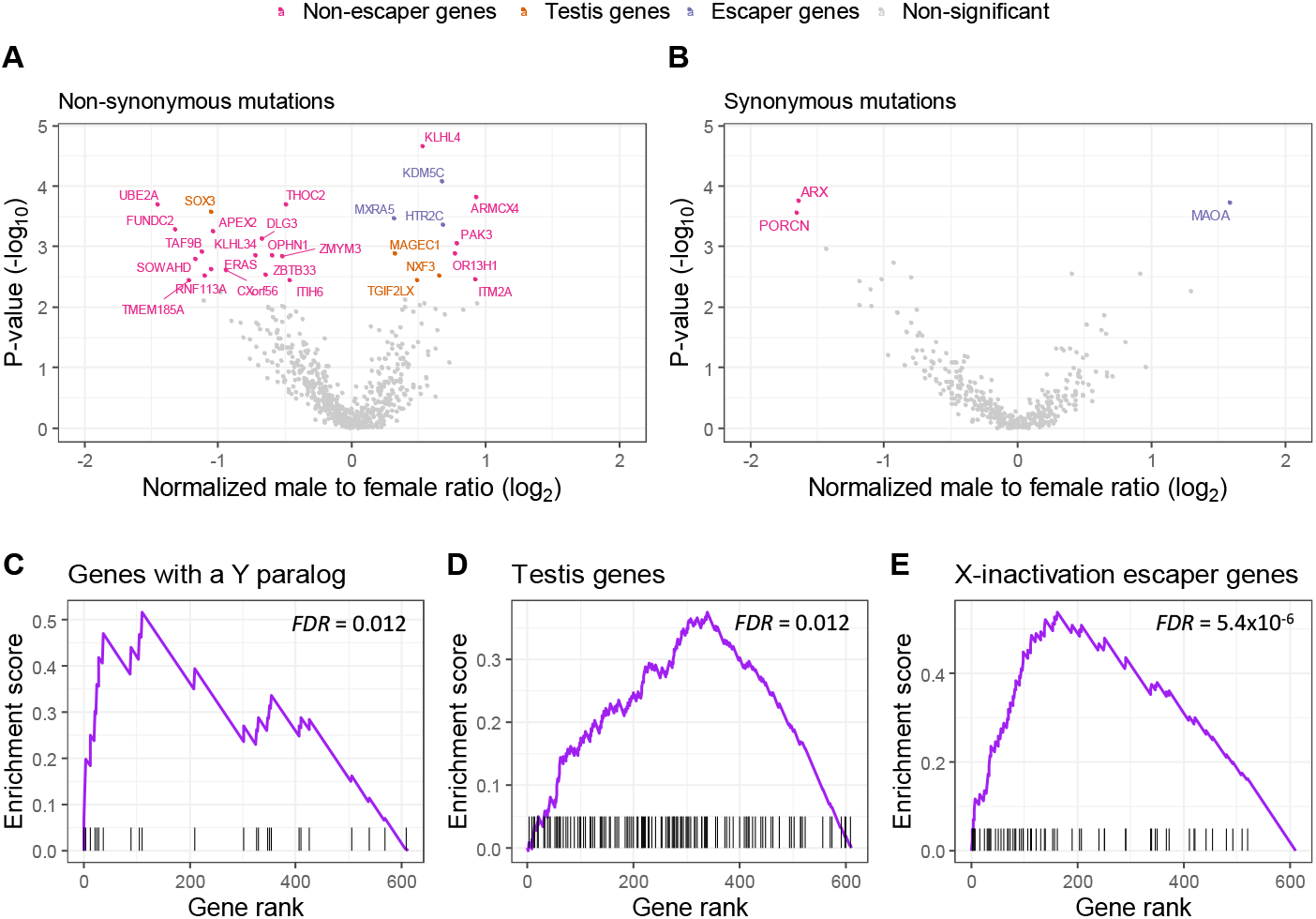
Sex differences in the rate of somatic mutations in human cancers for genes on the X-chromosome. (A-B) Analysis of somatic mutation rates in male and female tumors. (A) Nonsynonymous (B) and synonymous mutations. For each X-linked gene, significance is plotted as a function of the male to female ratio in mutation numbers. The ratio was normalized so that zero is the mean mutation ratio for all genes. Genes with a negative ratio value are with more mutations in females. The color of the gene labels indicates different characteristics of the gene. (C-E) Enrichment analysis plots for three gene sets: (C) genes that escape X inactivation (D) genes that express predominantly in male tissues, and (E) genes that have a paralog on the Y chromosome. In each graph, the vertical black lines indicate the position of genes in the sets relative to all X-linked genes that are ranked from the most significant male-biased genes to the most significant female-biased gene.

### Somatic mutation patterns in cancer resemble results from cell lines

Our analysis identified hallmarks of X-SMEGs and Y-SMEGs that could point to specific mechanisms. We determined if sex-biased genes in tumors share the same features. We tested the enrichment of three gene sets: (1) Genes with paralog on the Y chromosome. Because of male-specific redundancy, those genes are expected to accumulate more deleterious somatic mutations in males. (2) Genes predominantly expressed in male tissues (CTA genes). Those genes were found by us to be more essential to female XX_het_ cells and thus should have fewer mutations in females. (3) Genes that escape X-inactivation. While highly essential genes that escape X-inactivation are less essential to XX cells, previous work showed that in males, there is a higher rate of loss-of-function mutations in genes that are tumor suppressors that escape from X-inactivation (Dunford et al., 2017).

Consistent with these expectations, we found that genes with Y-linked paralogs were overrepresented among the male-biased genes (*FDR* = 0.012) (Figure 6C). Likewise, genes predominantly expressed in the testis were underrepresented among female-biased genes (*FDR* = 0.012) (Figure 6D). In agreement with a previous study (Dunford et al., 2017), genes that escape X-inactivation were more likely to be male-biased genes (*FDR* = 5.4×10^−6^) (Figure 6E). None of the tested gene sets were significantly enriched using a synonymous-based gene list (*P* > 0.1).

## Discussion

Multiple human disorders show sex differences in prevalence and symptoms, but the mechanisms remain unclear. Our study shows that the phenotypic outcome of loss-of-function mutations can be modified by the biological sex and the sex chromosomes. We found such effects for 178 differentially essential genes. In addition, we also identified 30 X-linked genes with a significantly different rate of somatic mutations in tumors between the sexes. Despite the difference in methodology, statistical power, and type of mutations, the essentiality analysis in cancer cell lines and somatic mutation in tumors both implicate genes that escape from X-inactivation, genes that express predominantly in testis (cancer-testis antigens), and Y-linked genes in sex-specific genetic effects.

The first notable group of genes implicated in sex-dependent essentiality includes X-linked genes that escape from X-inactivation. Those genes, which have two active copies in females and one in males, can contribute to sex differences in two ways depending on the effect caused by their loss (conferring advantage or disadvantage to the cells). We found that highly essential genes that escape X-inactivation are more sensitive to mutations in X0 cells. This can result from the higher probability of a complete knockout when there is only one copy of the gene. The finding aligns with the known protective role of having two X chromosomes in Mendelian disorders with X-linked inheritance (Snell and Turner, 2018). We also found that genes with excess somatic mutations in males are enriched for genes that escape X-inactivation. The three escape genes that show the most significant male bias (*KDM6A, DDX3X*, and *KDM5C*) were previously identified (Dunford et al., 2017). Among the 30 significant genes we identified, two additional ones escape X-inactivation and show significantly more mutations in male tumors (*MXRA5* and *HTR2C*). It was argued that the increase in loss-of-function mutations in males found in X-linked genes that escape inactivation is because those genes are tumor suppressors and complete knockout of the genes is more likely to occur in males with a single gene copy (Dunford et al., 2017). However, we found that escape genes under both positive and negative selection in the cancer cell lines showed a bias towards accumulating more somatic mutations in male tumors. In the cancer cell lines, three of the escape genes have on average a positive dependency score (*KDM6A* score = 0.18; *MXRA5* score = 0.11; *HTR2C* score = 0.047), which may suggest that loss-of-function mutations in those genes can cause increased proliferation (consistent with their role as tumor suppressors). The other two male-biased escape genes have negative scores (*DDX3X* score = -1.15; *KDM5C* score = -0.11), show intolerance to missense mutations in humans (Karczewski et al., 2020), and have a paralog on the Y chromosome, thus may be explained by a different mechanism.

The second notable group of genes we found to be involved in sex differences are genes expressed predominantly in the testis, also known as cancer-testis antigens. It is well established that cancer-testis antigens, which express primarily in the germ cells (both in the testis and fetal ovary), are reactivated in multiple cancer types (Fratta et al., 2011; Gordeeva, 2018). Although their role in cancer is not fully understood, there is considerable evidence that they can contribute to cancer cell proliferation and survival (Bhan et al., 2012; D’Arcy et al., 2014; Gordeeva, 2018; Gure et al., 2005; Sakurai et al., 2004; Shigematsu et al., 2010). X-linked cancer-testis antigens constitute 46% of all cancer-testis antigen genes (Liu, 2019), but their function in sex differences is still unknown. We found that cancer-testis antigens are more essential to XX heterozygote cells than X0 cells, but not to XX homozygote cells (which lack X-inactivation). Consistent with a higher selection against mutations in females, cancer-testis antigens also showed a significant excess of somatic nonsynonymous mutations in tumors from males compared to females. There are some indications for possible differences in the function of cancer-testis antigens in males and females. Two studies found that loss of X-inactivation in tumors is associated with increased expression of several cancer-testis antigens (Kang et al., 2015b, 2015a). However, in both cases, the X-inactivation loss was associated with global demethylation, which is known to independently increase the expression of cancer-testis antigens (Fratta et al., 2011; Gordeeva, 2018).

The third group of genes is influenced by the presence of a Y chromosome. Our analysis identified 122 genes whose essentiality is modified by the Y chromosome (Y-SMEGs) and thousands of genes that show Y-dependent differential expression. These results indicate that despite its small size, genes on the Y chromosome regulate the function and expression of multiple genes, many of them associated with cells proliferation and survival. The most promising Y-linked genes that might underline this variability in expression and essentiality are the eight genes that are known to be dosage-sensitive regulators of gene activity (*UTY, EIF1AY, ZFY, RPS4Y, KDM5D, DDX3Y, USP9Y*, and *TBL1Y*) (Bellott et al., 2014). The eight genes all have X chromosome paralogs that escape from inactivation. However, the fact that 40% of the differentially expressed genes between Y^+^ and Y-cells were also differentially expressed between XX females and XY males can be explained by two options. Either the X–Y gene pairs are not entirely functional redundant, or other Y-linked genes also contribute to the expression differences.

Our analysis revealed some of the mechanisms that might be responsible for the differential essentiality that is dependent on the Y chromosome. We showed that the most striking Y-dependent effects are explained by Y-linked genes that can compensate for the loss of an X-linked paralog. This might also explain the excess somatic mutations in X-linked genes with a Y paralog in male tumors. Among the four genes we found to be explained by Y-specific redundancy (*DDX3X, EIF1AX, RPS4X*, and *ZFX*), two (*DDX3X* and *RPS4X*) were previously reported to have redundant roles with their Y paralogs based on functional experiments (Venkataramanan et al., 2020; Watanabe et al., 1993). For all four genes, the average dependency score in Y^+^ cells was still substantially lower than 0, indicating that the loss of those X-linked genes causes decreased cell viability even with compensation by Y-linked genes. This is consistent with the dosage sensitivity of those genes, as was suggested before (Bellott et al., 2014).

In addition to the direct effect of Y chromosome genes on essentiality, our study also implies the indirect influence of Y-linked genes through modifying the expression of autosomal genes and their paralogs. The two autosomal genes highlighted in our study, *RPP25L* and *CHMP4B*, are characterized by variation in essentiality that is substantially explained by the expression levels of their autosomal paralogs (*RPP25* and *CHMP4C*). Still, this mechanism potentially explains many other Y-SMEGs. The presence of a Y chromosome was significantly associated with increased essentiality of *RPP25L* and *CHMP4B*. Our results suggest that this is because of the reduced expression of *RPP25* and *CHMP4C* in cells with a Y chromosome. These results further demonstrate that loss of the Y chromosome may dramatically affect the expression and essentiality of autosomal genes.

Our study has several limitations that are mainly a result of the use of data from cancer cell lines and tumors. First, cancer cells have multiple genomic alterations (Mani and Chinnaiyan, 2010) that might affect the results. Second, our analysis measures gene essentiality for the proliferation and survival of cancer cells in-vitro, and some of the findings might be more relevant to cancer cells. Despite those limitations, we believe that the large size of the datasets used and the inclusion of cells from multiple different lineages that are not expected to have identical genomic rearrangements can partially compensate for the above limitations. The replication of a few genes and the biological properties by studying sex differences in somatic mutation rates also supports the relevance of our findings in vivo. Furthermore, our results are likely relevant to other proliferating tissues and particularly to developing tissues that share features with cancer cells (Ma et al., 2010).

The results show that both the X and Y chromosomes have a global influence on gene expression and the essentiality of genes. We show that comparing cells with different compositions of sex chromosomes enables better discovery of differential expression and essentiality than comparing males and female cells. Our approach can be extended to other phenotypes, specific cell types, and developmental stages. In addition to the implications of our results to studying the differences between males and females, they are relevant to understanding specific disorders and genetic alterations. This includes syndromes with sex chromosome abnormalities like Turner syndrome and Klinefelter syndrome and the mosaic loss of the Y chromosome frequently observed in both cancer cells and during the normal aging process of male individuals (Guo et al., 2020). Our results also reflect on the importance of studying gene function considering the existence of sex-specific effects.

## Methods

### Cancer cell lines data acquisition and processing

Data on gene essentiality, gene expression, and relative copy number was obtained from the DepMap project (20Q3) (Tsherniak et al., 2017) (files: Achilles_gene_effect.csv, CCLE_expression.csv, CCLE_gene_cn.csv, sample_info.csv). Data on Y chromosome genes relative copy number was obtained from the cancer cell line encyclopedia (CCLE) web portal for Y chromosome genes (Ghandi et al., 2019) (https://portals.broadinstitute.org). Data on paralog genes were obtained using the biomaRt R package (Durinck et al., 2009). Genes that were labeled as having a Y paralog are genes with at least one Y-linked paralog. Genes that were labeled as having an autosome paralog had an autosome paralog but did not have a Y-linked paralog. A gene was defined as an X-inactivation escaper gene based on previously reported combined X chromosome inactivation status (Tukiainen et al., 2017).

### Differential essentiality analysis between male and female cells

Differential essentiality analysis between male and female cells was performed using the R package *limma* (functions *eBayes* and *lmFit*) (Core Development Team, 2020; Ritchie et al., 2015). The *limma* package performs differential expression analysis and borrows information across genes to better estimate the variance. A gene was defined as significantly differentially essential based on *FDR* corrected P < 0.1.

### Classification of sex chromosomes ploidy

Classification of cells based on the ploidy of the Y chromosome was based on the relative copy number of Y chromosome genes compared to all other genes and gene expression of Y chromosome genes. For each cell line, we calculated the mean Y chromosome relative copy number. We defined a threshold for classification based on the distribution of the relative copy number of the Y chromosome in female cells relative to males. In addition, we performed a PCA analysis on the expression of Y-linked genes and defined a threshold for classification based on the limits of the two main clusters. Cells outside the main clusters were excluded from further analysis (N = 37). Six female cells that were classified as Y^+^ were excluded from the analysis. Cells lacking copy number data were classified only based on the gene expression (N = 158).

Classification of the ploidy of the X chromosome was based on the relative copy number of X-linked genes and the level of heterozygosity. X chromosome heterozygosity was calculated based on SNP array data (downloaded from GEO Accession: GSE36138) (Barretina et al., 2012). SNPs were called using the *CRLMM* R package (Carvalho et al., 2010), and the percentage of heterozygote X chromosome SNPs was calculated for common (minor allele frequency > 10%) high confidence SNPs (mean confidence across samples > 0.95). The threshold for classification based on the X chromosome relative copy number was determined based on the location of the local minimum between the two peaks representing one or two copies of the X chromosome. Since the distribution for males and females was different, we used different thresholds. Similarly, we defined a threshold for classifying cells with heterozygote or homozygote X chromosomes that best separates cells as XX or X0 based on copy number data. Six female cells with conflicting data in the copy number and heterozygosity were excluded from the analysis. Female cells lacking SNP array data (N = 179) were classified based only on the relative copy number of X-linked genes.

### Estimation of the separate effects of the sex chromosomes and the biological sex on the essentiality

Two linear mixed-effect models were used to estimate the effect of the sex chromosomes and the biological sex on the 29 sex-dependent genes. In the first model, used to estimate the effect of the biological sex, the X and Y chromosomes were treated as random effects and the biological sex as a fixed effect. In the second model, used to estimate the effect of the X and Y chromosomes, the biological sex was treated as a random effect, and the X and Y chromosomes were defined as fixed effects. The models were tested with the *lme4* R package (Bates et al., 2015).

### Comparing X chromosome methylation levels between samples

Data on gene-wise methylation levels (promoter 1kb upstream TSS) was downloaded from the CCLE web portal (https://portals.broadinstitute.org) (Ghandi et al., 2019). Methylation levels were available for 651 samples (82%). The mean X chromosome methylation level was calculated for each sample based on 386 genes known to be inactivated (excluding escape genes). Comparison of mean methylation levels between XX heterozygote female cells vs. all other samples was performed using Welch’s two-sample t-test.

### Differential essentiality and differential expression analysis

Differential essentiality between XX_Het_ and X0 cells was performed similarly to the procedure used to compare male and female cells, but with the biological sex as an additional factor in the model. For X-linked genes that were significantly more essential to XX_Het_ cells, we compared the dependency score in XX_Het_ vs. XX_Hom_ females using Welsh’s t-test. Fisher’s exact test was used to test the enrichment of X-inactivation escaper genes in genes that are significantly more essential to X0 compared to XX_het_ cells. Enrichment of genes that are significantly more essential to XX_het_ cells than X0 cells in different tissues and identification of genes predominantly expressed in the testis was based on the Tissue-Specific Expression Analysis (TSEA) tool (Dougherty et al., 2010). Differential essentiality and differential expression between male cells Y^+^ cells vs. Y^-^ cells was performed using the limma R package with the biological sex and the number of X chromosomes as additional factors in the model.

### Mapping regions most likely to contain the causal gene for Y-SMEGs

The mapping of regions most likely to contain the causal gene for Y-SMEGs was based on cells with a partial deletion of the Y chromosome as indicated by the copy number of 42 Y-linked genes. Fourteen cells with partial Y chromosome deletion were used (with 10% to 85% of Y-linked genes present). The Y chromosome was divided into ten sections according to the overlap in the deletions. We calculated the difference in standard deviations between the observed and the expected dependency scores for each gene and section, assuming that the causal gene resides in the section. The section with the minimal difference between the expected and observed score was considered the most likely to contain the causal gene. We used Welch’s t-test for the difference in dependency score between cells with and without the section most likely to contain the causal gene and the score for Y^+^ and Y^-^ cells.

The difference between the observed and the expected dependency scores was calculated according to the following equation: 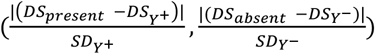, where the *DS*_*present*_ is the mean dependency score of cells with a deletion that does not include the section; the *DS*_*absent*_ is the mean dependency score of cells with a partial deletion which includes the section; *DS*_*Y+*_ is the mean dependency score for Y^+^ cells; the *DS*_*Y-*_ is the mean dependency score for Y^-^ cells; the *SD*_*Y+*_ is the standard deviation for the dependency score of Y^+^ cells; and the *SD*_*Y-*_ is the standard deviation for the dependency score for Y^-^ cells.

### Somatic mutations in tumors, data acquisition, and processing

Somatic mutations from whole-genome screens were obtained from the COSMIC (Catalogue of Somatic Mutations in Cancer) website (Tate et al., 2019). We excluded mutations in the mitochondria genome, non-PAR genes on the Y chromosome, and non-coding regions. We also removed mutations from unknown tissues and sex-specific organs (testis, prostate, placenta, ovary, breast, cervix, endometrium, genital tract, and penis). In addition, to avoid duplication in the data, we analyzed only mutations from canonical transcripts. We also kept only whole-genome screens and filtered out targeted studies. Moreover, we excluded samples with an outlier distribution of mutations, including a relatively low number of mutations on the X chromosome compared to the autosomes, and samples with a substantially low ratio between the number of unique mutated genes and the total mutations. The mutations were categorized as nonsynonymous (missense, nonsense, and frameshift) and synonymous. After applying all the filtrations, the dataset consisted of 1,336,600 nonsynonymous and 3,356,129 synonymous mutations from 7,489 males and 4,755 females.

### Comparison of somatic mutation rate between the sexes

We performed a randomization test to find genes with more mutation in one of the sexes across the different tumor types, similar to the method used in a previous study (Dunford et al., 2017). Separate tests were performed for synonymous and nonsynonymous mutations and the X chromosome and autosomes (including the PAR). The status of the genes was treated as binary, with or without a mutation. We only analyzed mutations from the 12 most common tissues in the database, which harbored ∼97% of the mutations. For each tissue and gene, the mutation probability in males was the number of mutations in males divided by the total number of mutations across all genes. The male probability for a mutation in each tissue was used to generate random numbers from a binomial distribution, summed across tissues, to have the expected number of mutations in males under the null hypothesis. This simulation was repeated one million times. The distribution was used to calculate a one-sided P-value based on the number of simulations where the number of male mutations was higher than observed, divided by the number of simulations. The P-values were transformed into a two-sided and corrected for multiple testing using the Benjamini-Hochberg FDR procedure. Genes with FDR corrected *P* < 0.1 were considered as significant. We used a gene set enrichment analysis approach to test for the enrichment of X-inactivation escaper genes, genes with paralog on the Y-chromosome, and genes that express predominantly in the testis. The enrichment was done on genes ranked by their *P*-value using the *fgsea* R package (Sergushichev, 2016).

