## Supplemental figures for "Gene essentiality in cancer cell lines is modified by the sex chromosomes"

Supplemental Information:

Figure S1. The essentially of CDK6 and PAX8 is influenced by female-only tissues but not by the sex of the cells.

Figure S2. Identification of Y chromosome ploidy and the effect of X chromosome ploidy on X-inactivation.

Figure S3. Testis-specific genes are essential for $XX_{het}$ cells but not for $XX_{hom}$ cells.

Figure S4. Prediction of the locations on the Y chromosome that contains the causal gene for Y-SMEGs.

Figure S5. A specific region on the Y chromosome modifies essentiality.

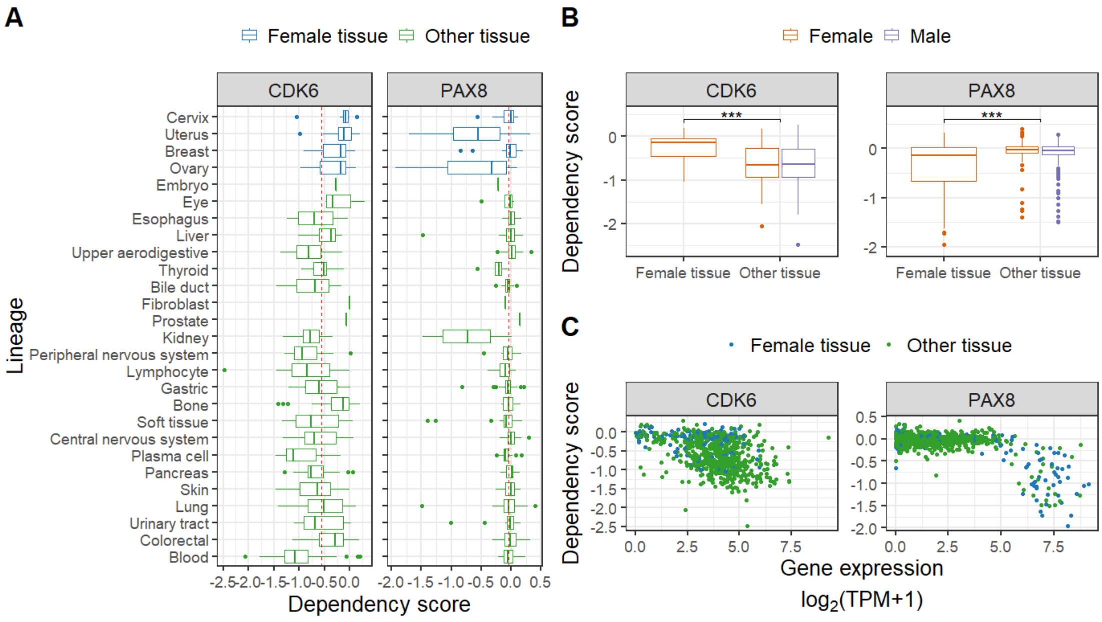


**Figure S1. The essentially of CDK6 and PAX8 is influenced by female-only tissues but not by the sex of the cells.**

(A) The dependency of CDK6 and PAX8 according to the lineage of the cell. Female-only tissues are in blue, and all other tissues are in green. The dashed line is the median dependency for all cells.

(B) The dependency of CDK6 and PAX8 in female-only tissues and all other tissues. Female cells are in orange and male cells in purple. ***, *P* <0.001

(C) Relationship between the essentiality (dependency score) and expression for CDK6 and PAX8. Female tissues are in blue, and all other tissues are in green.

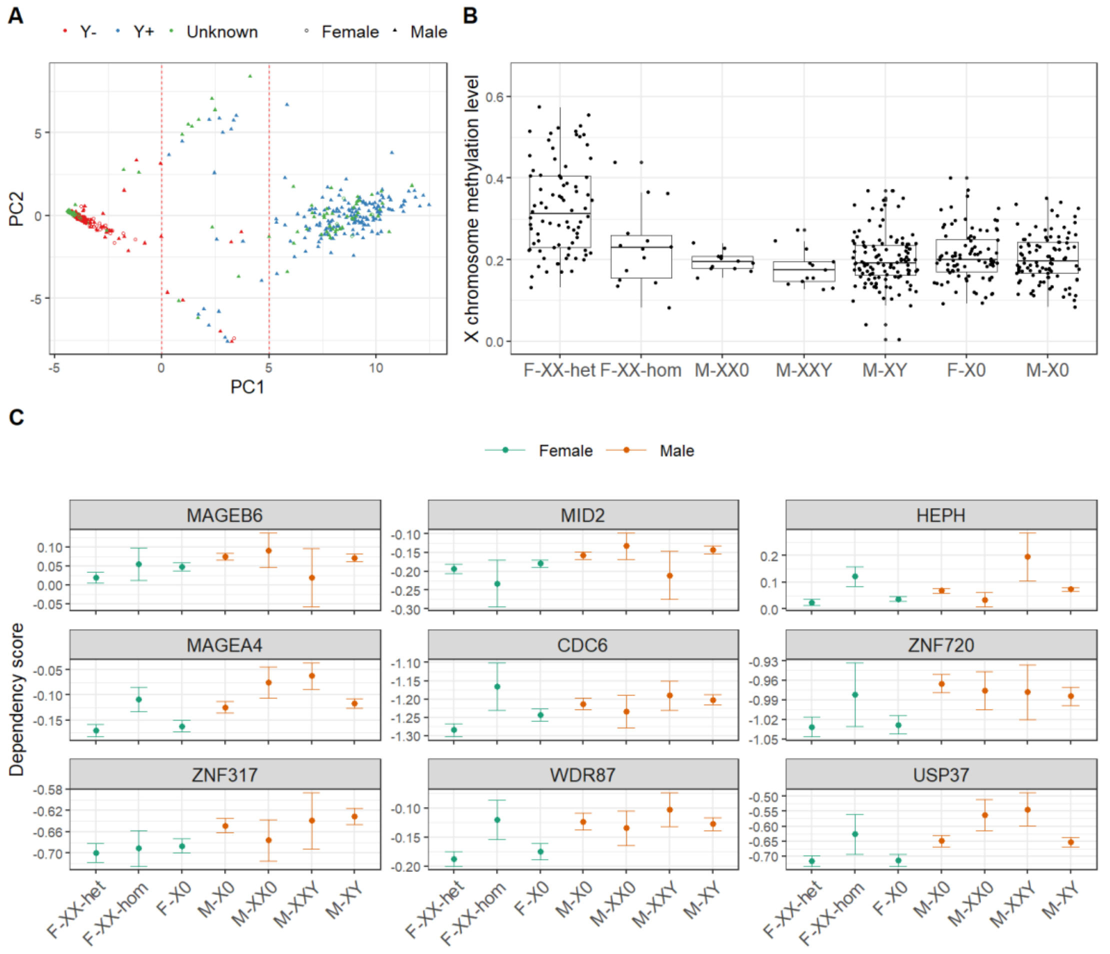


**Figure S2. Identification of Y chromosome ploidy and the effect of X chromosome ploidy on X-inactivation.**

(A) A PCA plot of the expression of Y chromosome genes. Cells with PC1 > 5 were defined as $Y^+$, cells with PC1 < 0 were defined as $Y^-$, and cells with 0 < PC1 < 5 were excluded from further analysis. Dashed lines indicate the threshold for definition as $Y^+$ and $Y^-$ cells. The color is the cell classification based on the relative copy number of the Y chromosome.

(B) Mean percentage of methylated CpGs in the promotors of X chromosome genes that do not escape X-inactivation. Cells are divided by their sex chromosome ploidy. The difference between females $XX_{Het}$ and all other samples is highly significant ($P$ = $2.6x10^{-15}$).

(C) Genes with a significant difference in essentiality between male and female cells not affected by the sex chromosomes (additional genes, not shown in Figure 1F). Values are the essentiality (mean dependency score) per genotype ± standard error of the mean (SEM).

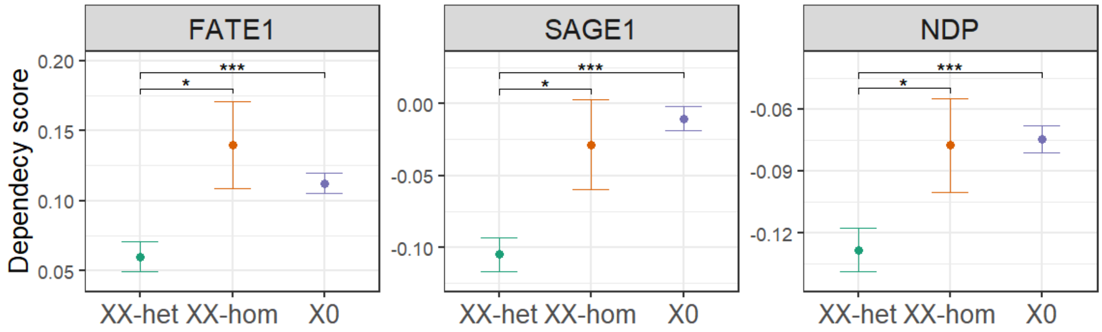


**Figure S3. Testis-specific genes are essential for $XX_{het}$ cells but not for $XX_{hom}$ cells**.

X-linked genes that are significantly more essential to $XX_{Het}$ cells than X0 and $XX_{hom}$ cells (additional genes not shown in Figure 2D). Values are mean dependency score ± SEM. * *P*<0.05; **, *P*<0.01; ***, *P*<0.001.

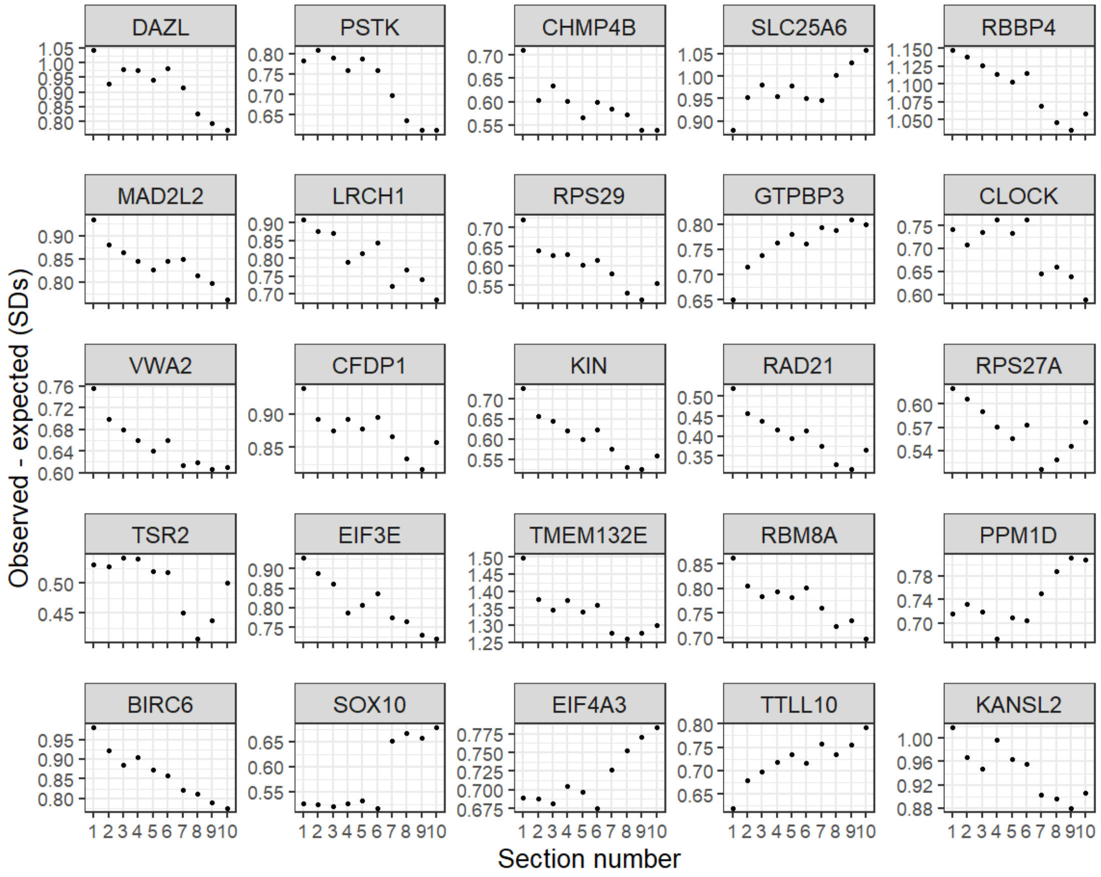


**Figure S4. Prediction of the locations on the Y chromosome that contains the causal gene for Y-SMEGs.** The difference (in standard deviations) between the expected and observed dependency scores assuming the causal gene is in the indicated area.

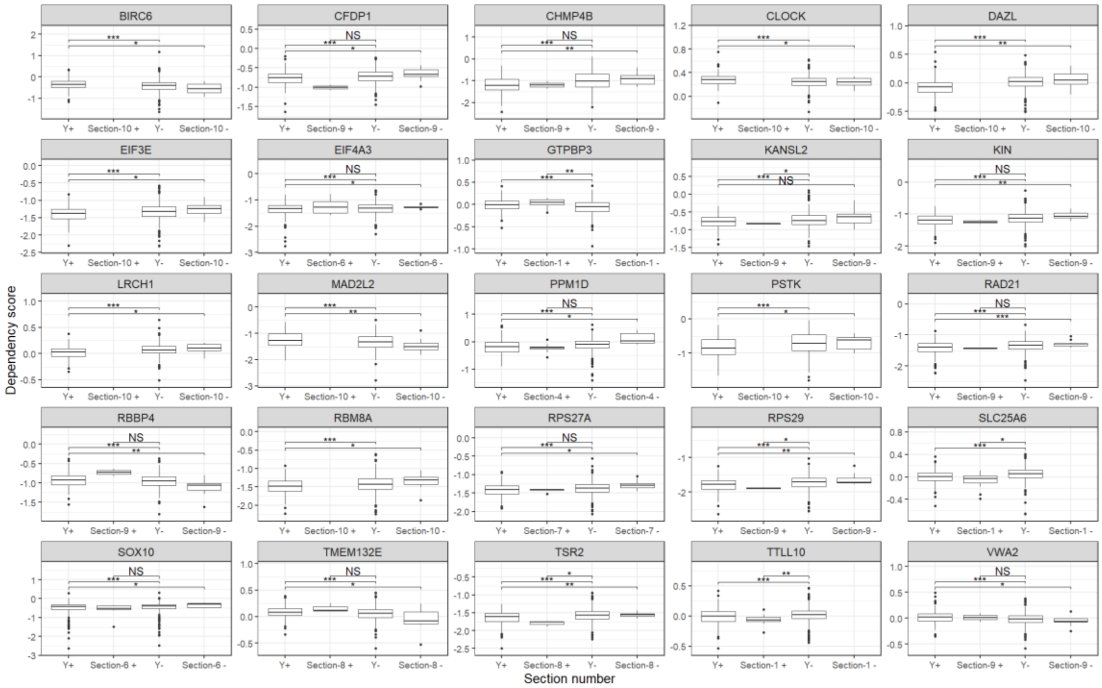


**Figure S5. A specific region on the Y chromosome modifies essentiality.**

The boxplots show the essentiality (dependency score) for cells with and without the section most likely to contain the causal gene and $Y^{+}$ and $Y^{-}$ cells. NS, $P > 0.05$; *, $P<0.05$; **, $P<0.01$; ***, $P<0.001$.
